# Drug combination sensitivity scoring facilitates the discovery of synergistic and efficacious drug combinations in cancer

**DOI:** 10.1101/512244

**Authors:** Alina Malyutina, Muntasir Mamun Majumder, Wenyu Wang, Alberto Pessia, Caroline A. Heckman, Jing Tang

## Abstract

High-throughput drug sensitivity screening has been utilized for facilitating the discovery of drug combinations in cancer. Many existing studies adopted a dose-response matrix design, aiming for the characterization of drug combination sensitivity and synergy. However, there is lack of consensus on the definition of sensitivity and synergy, leading to the use of different mathematical models that do not necessarily agree with each other. We proposed a cross design to enable a more cost-effective testing of sensitivity and synergy for a drug pair. We developed a drug combination sensitivity score (CSS) to summarize the drug combination dose-response curves. Using a high-throughput drug combination dataset, we showed that the CSS is highly reproducible among the replicates. With machine learning approaches such as Elastic Net, Random Forests and Support Vector Machines, the CSS can also be predicted with high accuracy. Furthermore, we defined a synergy score based on the difference between the drug combination and the single drug dose-response curves. We showed that the CSS-based synergy score is able to detect true synergistic and antagonistic drug combinations. The cross drug combination design coupled with the CSS scoring facilitated the evaluation of drug combination sensitivity and synergy using the same scale, with minimal experimental material that is required. Our approach could be utilized as an efficient pipeline for improving the discovery rate in high-throughput drug combination screening. The R scripts for calculating and predicting CSS are available at https://github.com/amalyutina/CSS.

**Author summary:** Being a complex disease, cancer is one of the main death causes worldwide. Although new treatment strategies have been achieved with cancers, they still have limited efficacy. Even when there is an initial treatment response, cancer cells can develop drug resistance thus cause disease recurrence. To achieve more effective and safe therapies to treat cancer, patients critically need multi-targeted drug combinations that will kill cancer cells at reduced dosages and thereby avoid side effects that are often associated with the standard treatment. However, the increasing number of possible drug combinations makes a pure experimental approach unfeasible, even with automated drug screening instruments. Therefore, we have proposed a new experimental set up to get the drug combination sensitivity data cost-efficiently and developed a score to quantify the efficiency of the drug combination, called drug combination sensitivity score (CSS). Using public datasets, we have shown that the CSS robustness and its highly predictive nature with an accuracy comparable to the experimental replicates. We have also defined a CSS-based synergy score as a metric of drug interaction and justified its relevance. Thus, we expect the proposed computational techniques to be easily applicable and beneficial in the field of drug combination discovery.

## Introduction

Despite the great advances in the understanding of cancer, there remains a major gap between the vast knowledge of molecular biology and effective anti-cancer treatments. Next generation sequencing has revealed the intrinsic heterogeneity in cancer survival pathways, which partly explains why patients respond differently to the same therapy [1]. To reach effective and durable clinical responses, cancer patients who become resistant to standard treatments need multi-targeted drug combinations, which shall effectively inhibit the cancer cells and block the emergence of drug resistance [2–4].

In order to predict novel drug combinations, high-throughput drug screening has been applied on a large variety of cancer cell lines and more recently on patient-derived cancer samples [5–6]. Ideally, a promising drug combination should achieve the therapeutic efficacy at reduced dosages, and therefore also minimize the toxicity and other side effects associated with high doses of single drugs [7–8]. Therefore, both the sensitivity and synergy of a drug combination need to be considered when evaluating the high-throughput screening results for further validation [9].

Many high-throughput drug combination screens combine two drugs at a full matrix of multiple doses, for which the cell viability or growth inhibition effects are measured [10–11]. The drug combinations are usually ranked based on the degree of synergy, such that the drug combinations that produce higher growth inhibition effects compared to the single drugs will be prioritized. However, there have been multiple methods to score drug synergy, each of which relies on a different mathematical model that do not fully agree with each other [12]. The lack of consensus on the choice of synergy scoring methods may partly explain the difficulty to validate drug combination discoveries in a high-throughput setting [13]. On the other hand, focusing only on synergy but not on sensitivity may produce false positive drug combinations that do not necessarily reach therapeutic efficacy, despite being synergistic [14]. However, unlike the sensitivity of single drugs which can be directly derived from dose-response curves [15], the sensitivity of a drug combination remains largely undefined, as the same sensitivity can be achieved using different dose combinations. Furthermore, there is a lack of scoring approaches to fully capture the synergy and sensitivity simultaneously.

On the other hand, the dose-response matrix utilizes a full factorial design to test multiple dose combinations, and thus demands a relatively large amount of cancer cells. For patient-derived cancer samples which are typically rare and restricted in volume, the full dose-response matrix design may be infeasible for testing even a minimal number drug combinations. Furthermore, cancer samples of different genetic profiles are known to respond differently to the same drug combination. With the limited amount of drug combinations as the training data, it becomes a daunting task for any machine learning approach to navigate the combinatorial space to pinpoint the most promising drug combinations that are specific for individual cancer samples [16].

To overcome these challenges, we proposed a cost-effective experimental and computational procedure to facilitate the prediction of drug combination synergy and sensitivity. We utilized an experimental design where two drugs are crossed at their IC_50_ concentrations, and either drug is allowed to span over multiple doses while the concentration of the other drug is fixed. The resulting dose-response curves are utilized to defined a drug combination sensitivity score (CSS). Using a large scale of drug combination study, referred to as the O’Neil data [17], we showed that the CSS is highly reproducible, suggesting its robustness as a metric for characterizing drug combination responses. Furthermore, we found that the CSS can be predicted at high accuracy using chemical and pharmacological features of the drug combinations. Based on the difference between the observed and expected CSS values, a drug synergy score can be determined straightforwardly. We showed that such a CSS-based synergy score can also detect the true synergistic and antagonistic drug combinations with high accuracy. Compared to the dose-response matrix design, the cross design requires minimal amount of experimental materials, while it still maintains a sufficient level of accuracy for capturing both synergy and sensitivity simultaneously. We foresee that such an experimental design and its CSS scoring would facilitate the standardization of drug combination analysis that is currently lacking in functional chemical screening, and would allow for the scale up of drug combination testing eventually for personalized medicine.

## Results

### CSS values are highly reproducible and robust

We applied the CSS scoring on the O’Neil drug combination data, which consists of 22,737 drug combinations for 39 cancer cells [17]. We found that the CSS_1_ and CSS_2_ values calculated using either drug fixed at its IC_50_ concentration are highly correlated (Pearson correlation = 0.82, p-value = 2×10^−16^; Fig 1A). Both CSS_1_ and CSS_2_ values ranged from 0 to 50, with a marginal absolute difference of 5.62 (Fig 1B). Such a high level of consistency holds true for all the 39 cancer cell lines and the majority of the 38 unique drugs, suggesting the robustness of the CSS scoring method (Figs 1C and D, Fig S1). We also found a high correlation between the CSS value and those derived from single replicates (minimal correlation = 0.97, Table S3). In order to check whether the CSS values are within the range of the CSS replicates for each drug combination, we calculated the minimal and maximal values over the CSS replicates for each drug combination and plotted them together with CSS values over the standard deviation of the CSS replicates. For a better visualization, we applied a generalized additive model to smoothen the CSS lines and obtain 95% pointwise confidence interval around the mean (Fig S2). Only 4% of the drug combinations have the CSS values being out of the CSS replicate-based limits, however this can be explained by the higher variance over the replicates.

**Fig 1.**
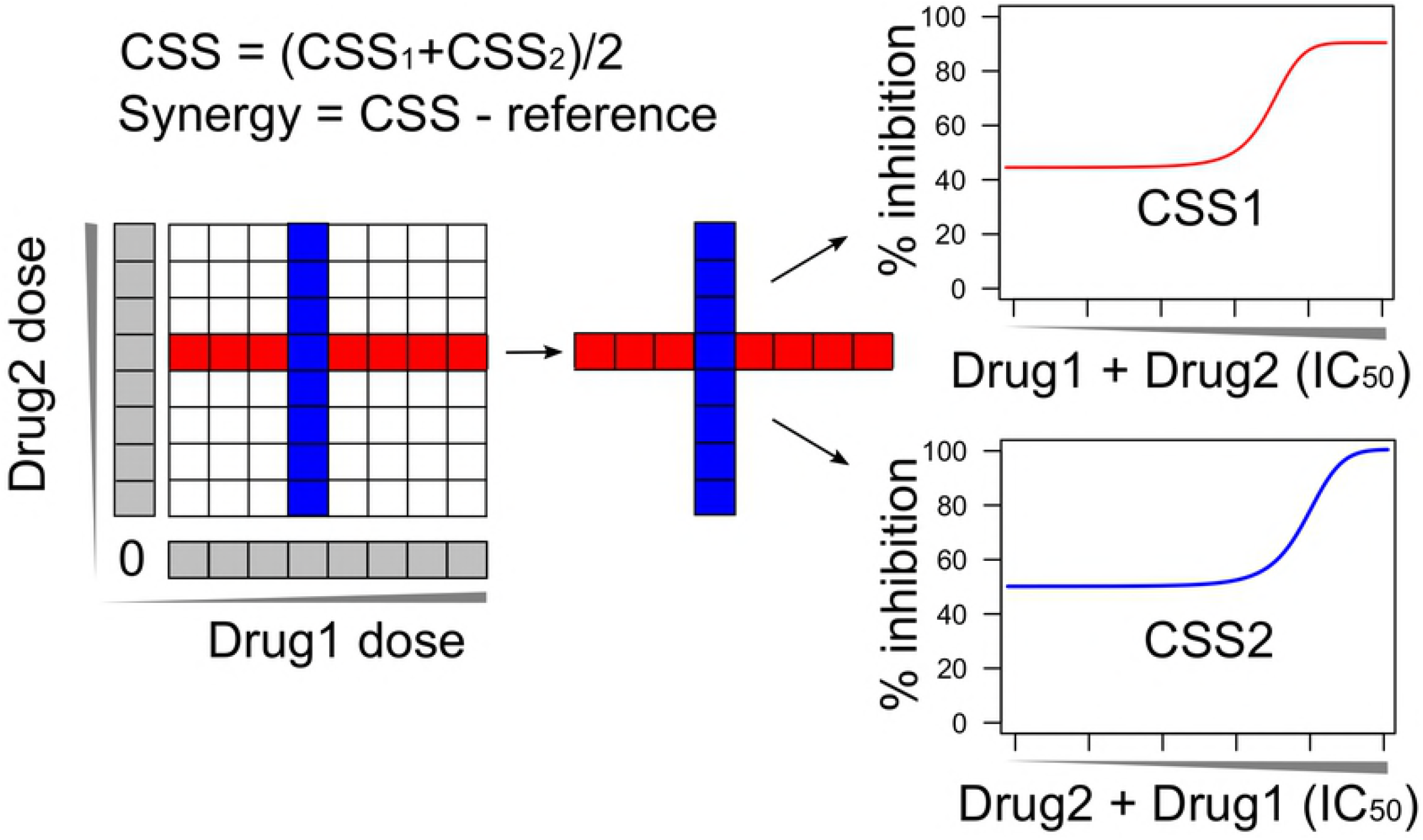
Robustness and replicability of CSS. (A) The correlation of CSS_1_ and CSS_2_ over all the drug combinations; (B) Density plot of the CSS_1_ and CSS_2_ distributions; (C) The correlation per cell line colored according to the tissue type; and (D) The correlation per drug colored according to the drug target class.

Notably, we found that drug combinations that involved bortezomib showed much lower correlation (0.26) between the CSS_1_ and CSS_2_ values compared to other drug combinations. Since the O’Neil data contains the replicates for single drug screening, we analyzed the coefficient of variation (CV) of the cell viability readout for each drug in the replicates. As expected, we found that bortezomib has the highest CV (0.28), suggesting a relative low quality of the drug sensitivity data involving this drug (Fig S3). Therefore, the lower correlation between the CSS_1_ and CSS_2_ for a drug combination may be attributed to a higher experimental variation, and thus were considered as low quality data points. For the subsequent analysis, we selected only those drug combinations that have the absolute difference between CSS_1_ and CSS_2_ lower than 10, resulting in a total of 18,905 drug combinations. After this filtering the correlation between CSS_1_ and CSS_2_ was further improved (Pearson correlation = 0.93, p-value = 2×10^−16^). Furthermore, the mean absolute difference between CSS_1_ and CSS_2_ was 3.83, which became comparable to the variability determined from the technical replicates of CSS_1_ and CSS_2_ (2.92 and 3.06 respectively), suggesting that the difference between CSS_1_ and CSS_2_ is similar to what is expected when repeating the experiment. Taken together, CSS_1_ and CSS_2_ values are highly consistent and therefore supported their averaging as a summary for the drug combination sensitivity score.

### CSS can be predicted using machine learning approaches

Given that the CSS is highly reproducible as a summary of the overall sensitivity of a drug combination, we explored whether CSS can be predicted using pharmacological and chemical information of the drugs. We considered a drug combination as a combination of its drugs target profiles as well as their chemical fingerprints, with which the machine learning approaches illustrated in the previous section can be optimized by exploring the feature space using the training data. We examined three major machine learning methods for predictions: Elastic Net, Random Forests and Support Vector Machines.

We found that all of these machine learning approaches worked reasonably well, with the Elastic Net consistently achieving the best performance, with the mean MAE of 4.01 which is comparable to that (2.07) of a technical replicate (Table 1). Note that in our cross-validation setting the drug combinations in the test data were not present in the training data, the machine learning methods were still able to predict the CSS values for new drug combinations by exploring the feature similarity in the drug targets and chemical fingerprints. The prediction performance thus validated our hypothesis that a drug combination can be considered as a combination of their drug target profiles and chemical-structural properties, with which the CSS score can be predicted with high confidence using the state-of-the-art machine learning approaches.

**Table 1.** The prediction performance for Elastic Net, Random Forests and Support Vector Machines, as compared to the upper limit when selecting randomly one technical replicate as the prediction.

| *Method* | *RMSE* | *R2* | *COR* | *MAE* |
| --- | --- | --- | --- | --- |
| *Elastic Net* | 5.20±1.11 | 0.80±0.06 | 0.90±0.03 | 4.01±0.86 |
| *Random Forests* | 6.30±1.18 | 0.71±0.07 | 0.85±0.04 | 4.75±0.9 |
| *Support Vector Machines* | 7.47±1.32 | 0.57±0.08 | 0.80±0.04 | 5.80±1.07 |
| *Technical replicate* | 2.87±0.59 | 0.93±0.04 | 0.97±0.02 | 2.07±0.45 |

All the values are mean+/-standard deviation.

Since both the drug target profiles and chemical fingerprints were considered as the drug combination features, we next evaluated their prediction performances separately using the Elastic Net method. For drug-target profiles we collected known targets that were experimentally validated as well as the additional secondary targets that were predicted with high confidence using the SEA method. For chemical fingerprints we used the MACCS fingerprint which contains 166 structural features [18]. As expected, when combining all the features the model achieved the best performance (Table 2). We found that in general drug target profiles were predictive of CSS, especially when including the experimentally validated targets. The predicted targets using the SEA method did not improve the prediction accuracy significantly, indicating that even though secondary drug target interactions may occur, most likely they have minor functional impact that may not lead to the changes in cancer cell viability and thus does not contribute to the prediction of CSS. On the other hand, we found that chemical fingerprints were less predictive of CSS compared to the drug-target profiles, suggesting that the use of MACCS might be suboptimal to capture the relevant structural information for predicting the drug combination sensitivity. However, as the focus of this study was to show the validity of using machine learning methods to predict the CSS score, we decided to explore other chemical fingerprint features as a future step.

**Table 2.**
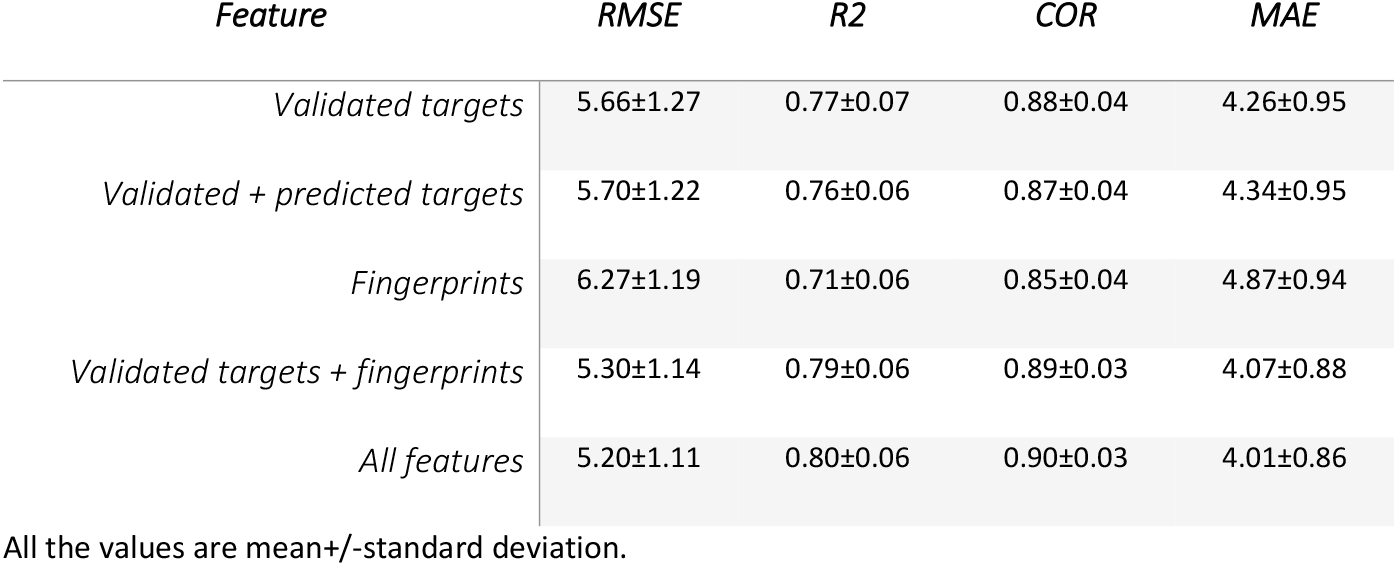
The prediction performances for drug-target features and chemical fingerprint features using Elastic Net.

We considered the regression coefficients that were determined in the Elastic Net model as an indication of their importance to contribute to the CSS prediction. We collected 67 features that have their absolute coefficients greater than 3 for at least one cell line. Unsupervised hierarchical clustering with the Manhattan metric was then applied to group the cell lines and the selected features (Fig 2). We found that certain drug target features were present with high coefficients across all the cell lines. For example, DNA topoisomerases including TOP1MT, TOP2A, TOP2B and TOP1 were selected, with an average coefficients of 8.2, 2.7, 2.6 and 1.0 separately. Despite the difference in the level of variable importance, all the DNA topoisomerases showed positive coefficients in 38 of 39 cell lines, suggesting that targeting DNA topoisomerases were associated with a higher CSS. DNA topoisomerases are known proteins which are essential for cell replication and metabolism [19]. Including a topoisomerase inhibitor can thus enhance the drug combination sensitivity in many cancer cell lines. On the other hand, the only cell line that showed negative coefficients for TOP1MT was LNCAP (prostate cancer), which turned out to be the cell line that has the smallest average CSS scores for drug combinations involving the TOP1MT inhibitor (topotecan) (Fig S4).

**Fig 2.**
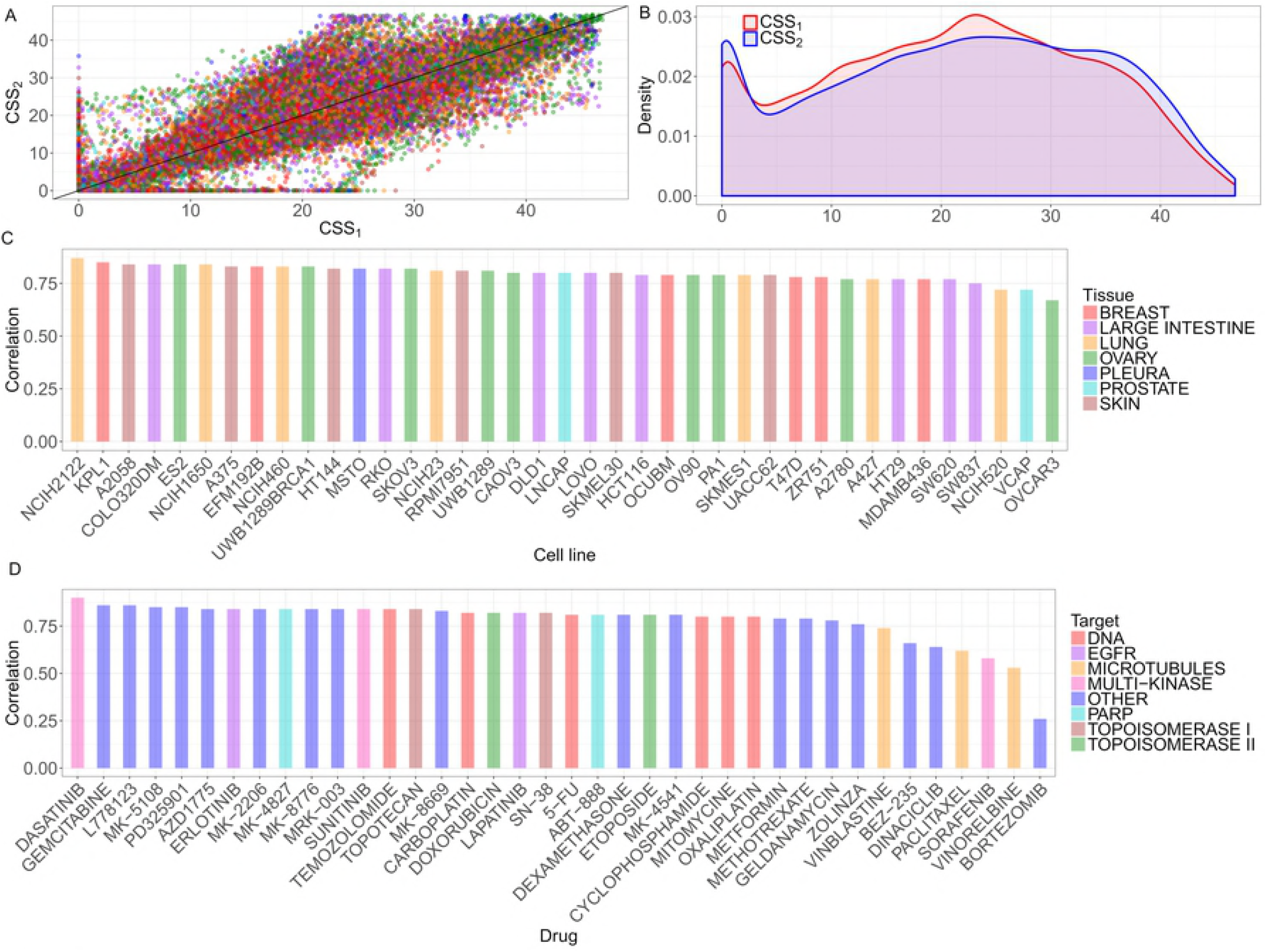
The feature importance map by Elastic Net for each cell line. Cell-line independent as well as cancer subtype-specific features can be identified by evaluating the regression coefficients of the Elastic Net model. Features such as TOP1MT, TOP1, TOP2A/B has shown consistently positive coefficients as compared to features such as AKT1/2/3 which showed cancer subtype specificity in breast cancer (indicated as arrows).

At the cell line level, we found that cell lines of the same tissue type did not necessarily cluster together, indicating their distinctive drug combination response profiles. However, we found that the breast cancer cell lines did form a major cluster including KPL1, ZR751, EFM192B, OCUBM and T47D, while the only outlier was MDAMB436. Indeed, MDAMB436 is the only triple negative breast cancer (TNBC) subtype, while the other cell lines are either ER positive (KPL1, ZR751 and T47D), or HER2 positive (EFM192B and OCUBM). It has been known that TNBC respond anticancer drugs differently from ER and HER2 positive breast cancers due to the distinctive disease mechanisms [20]. The features selected for the CSS prediction separated these two distinctive breast cancer subtypes, suggesting the validity of using CSS to cluster cancer of different subtypes. Furthermore, we found that AKT targets (AKT1/2/3) were among the top ones that showed higher importance in the non-TNBC group. A combination of an AKT inhibitor and TOP1MT inhibitor therefore can be suggested to treat non-TNBC but not necessarily for TNBC breast cancers. On the other hand, we found that CHEK1 and PARP3/4 targets were selected for MDAMB436 but not for non-TNBC group, suggesting that a combination of a CHEK inhibitor and PARP inhibitor might be tested for TNBC, but not for non-TNBC. The mechanisms of actions for the proposed drug combination may prove interesting for experimental validations. Taken together, the features that were selected from the CSS prediction may help to pinpoint the underlying target interactions, which are of pivotal importance to identify the drug combination response biomarkers.

### The CSS-based synergy scores can predict the true synergy and antagonism

Next, we defined the degree of drug synergy as the differences between the dose-response curves of a drug combination and its single drugs. We derived three variants of the CSS-based synergy score (S_sum_, S_max_, S_mean_) and compared them with the HSA, Bliss, Loewe and ZIP synergy scores that were determined using the full-dose response matrix. We evaluated whether the CSS-based synergy scores using the cross design can capture the ground truth. Although determined using only one row and one column from the dose-response matrix, all the CSS-based synergy scores managed to obtain a good correlation with the synergy scores based on the dose-response matrix design (Table 3).

**Table 3.**
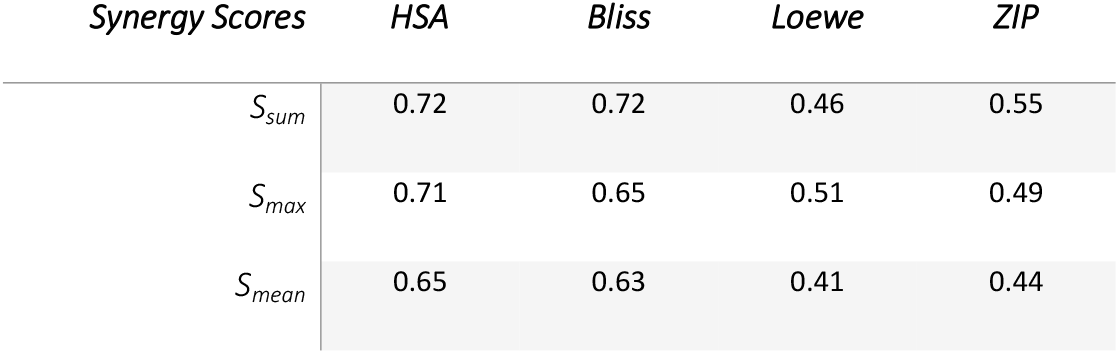
Correlations of the CSS-based synergy scores with those derived using four reference models that were calculated using the full dose-response matrices.

We found that the CSS-based synergy scores correlated relatively well with the HSA and Bliss scores, while the correlation started to decrease when comparing to the Loewe and ZIP scores. Since all the synergy scoring models utilized different assumptions for the reference of no synergy, we therefore did not expect a perfect correlations in such pairwise comparisons. Of all the three variations of CSS-based synergy score, we found that S_sum_ showed the best correlation with those determined using the full dose-response matrix. As S_sum_ considers the additive effect of single drug sensitivities as the expectation of no synergy, it thus can be considered a more conservative scoring method compared to S_max_ and S_mean_, where the maximal and average effect of single drugs were considered as reference separately. To control the false discovery rate of detecting synergistic combinations, we therefore proposed S_sum_ as a more appropriate scoring method for the cross drug combination design.

Furthermore, we evaluated the predicting accuracy of the CSS-based synergy scores for true synergistic and antagonistic drug combinations. We applied a stringent criteria to determine the ground truth from the dose response matrix data, such that all the synergy scores (HSA, Bliss, Loewe and ZIP) must be higher than 5, or lower than −5, to be classified as a true synergistic or antagonistic drug combination, respectively. From the O’Neil data, we identified 3,716 true synergistic and 218 true antagonistic drug combinations. We then asked the question of whether the CSS-based synergy scores that were determined using the cross design can predict the ground truth. We showed that the CSS-based synergy scores managed to achieve the area under the ROC curves of 0.997 (S_sum_), 0.996 (S_max_) and 0.992 (S_mean_) to detect the true synergistic and antagonistic combinations correctly (Fig 3A). Note that in order to calculate the synergy score, only two vectors of the drug combination responses are needed, rather than the full dose-response matrix. Therefore, the CSS-based synergy score needs a substantially fewer measurements compared to the other well-established synergy scores. Still, the CSS-based synergy scores can predict the most synergistic and antagonistic drug combinations with high accuracy. On the other hand, the CSS-based synergy score and the CSS drug combination sensitivity score were using the same unit as the percentage of the actual drug response compared to the theoretical upper limit. Therefore, the synergy score can be interpreted as the extra benefit of combining two drugs that can achieve an effect closer to the upper limit. We summarized both CSS drug sensitivity scores and CSS-based synergy scores for all the drug combinations as an S (sensitivity)-S (synergy) plot (Fig 3B; Table S4). By applying a threshold of the 3^rd^ quantiles for CSS and S, we can clearly identify the most promising drug combinations that fulfill both the sensitivity and synergy criteria, while avoiding the false positive drug combinations that might be synergistic but do not achieve a sufficient high level of sensitivity. Taken together, the combined use of CSS drug combination sensitivity score and its associated synergy score allows a simultaneous evaluation of the sensitivity and synergy for a drug combination, which will facilitate a more systematic analysis of high-throughput drug combination data with much less experimental materials.

**Fig 3.**
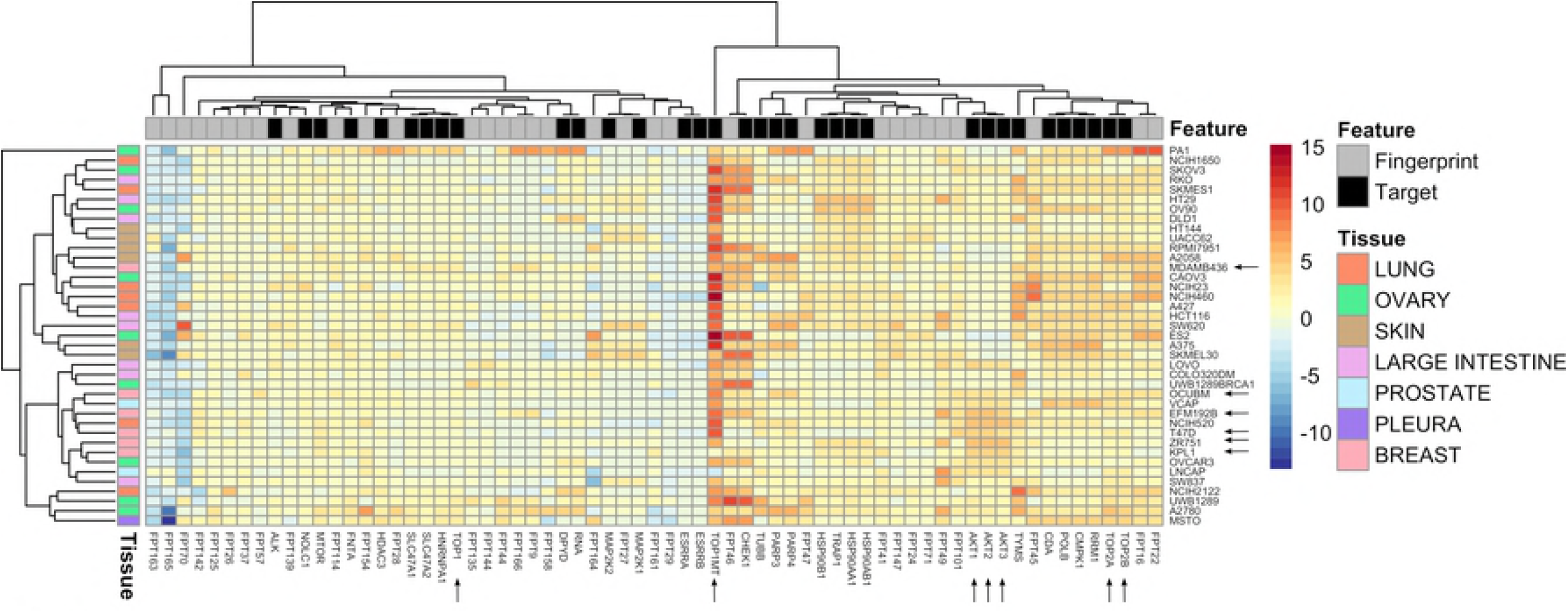
(A) The ROC curves for the CSS-based synergy scores to detect true synergistic and antagonistic drug combinations. (B) The S-S plot for all the drug combinations. The drug combinations with the 75^th^ percentile and above for both the CSS and the S scores were considered as the prioritized hits for further experimental validation and highlighted in red.

## Discussion

Drug combinations may potentially lead to more durable clinical responses by overcoming intra-tumoral heterogeneity and drug resistance to monotherapies. Identifying drug combinations that are tailored for personalized medicine is a challenge, as the number of possible combinations may easily grow exponentially [21]. High-throughput drug combination screening has been increasingly utilized for early detection of true synergistic and effective drug combinations. However, systematic identification of drug combinations is difficult, as the concepts of synergistic versus effective drug combinations are often intertwined and sometimes interchanged without sufficient clarification. Furthermore, there is a lack of consensus on what the definition of synergy is, which might contribute to the poor reproducibility of many drug combination studies. The uncertainty about the endpoint measurement in drug combination screens brings additional complexity for any machine learning approach to tackle the prediction problem.

We developed a novel scoring approach called CSS for drug combinations that can be efficiently determined using a simple experimental design. We found that the CSS is highly replicable and therefore can be considered as a robust metric to characterize drug combination sensitivity. To leverage the drug combination CSS data, we also developed a testing platform to allow for a systematic evaluation of the prediction accuracy of different machine learning methods. We found that the target information for the compounds as well as their chemical fingerprints are highly predictive of the CSS values, with an accuracy comparable to the experimental replicates. Therefore, the rationale of considering a drug combination as a combination of their target and fingerprint profiles can be justified. This would also allow the augmentation of single-drug screening and drug combination screening data to train a machine learning model, as many drugs are multi-targeted which are equivalent to a drug combination with the same target profile. In this study we focused on drug combination prediction within the same cell line. In the future, we would include the molecular features of the cell lines to improve the prediction accuracy as well as identify drug combination specific biomarkers across different cell lines. On the other hand, as the focus of the current study is to propose the new experimental design and to justify its associated drug combination scoring methods, we tested the predictability of CSS using conventional machine learning methods, and showed that CSS can be accurately inferred from the pharmacological features of drug combinations. More advanced machine learning methods such as Deep Learning [16] or network-based methods [22] may further improve the prediction accuracy, which will be tested as future work.

A truly promising drug combinations shall reach therapeutic efficacy via a strong synergy. While there have been multiple synergy scoring methods that can be applied to the full dose-response matrix design, they do not always produce the consistent results. The truly synergistic and antagonistic drug combinations may therefore be determined by finding the consensus across the different scoring methods [12]. We developed a CSS-based synergy score to quantify the degree of interactions in a drug pair, and showed that the CSS-based synergy score can identify the truly synergistic and antagonistic drug combinations accurately. Therefore, the CSS-based synergy score can be used for the prioritization of a primary drug combination screen using the cross design, after which only the significant drug combinations warrant a confirmation screen using the full dose-response matrix design. Furthermore, we proposed a novel S-S plot to visualize drug combination sensitivity and synergy using the same scale, which enables an unbiased way to explore high-throughput drug combination data more efficiently. On the other hand, the CSS is defined at the IC_50_ concentrations of the background drugs. Therefore, a synergistic drug combination determined by the CSS-based synergy score should be more therapeutically relevant than a drug combination where the synergy is detected at a higher concentrations, which are often associated with unwanted off-target effects and side-effects.

The advantage of the proposed cross design is that drug combination screens and single-drug screens can be implemented in a sequential manner, which requires much less cells and compound materials compared to a full dose-response matrix. With the introduction of CSS scoring, we are foreseeing a lower technical barrier to carry out large scale drug combination studies with minimal cellular materials. Although the drug combination data we explored here involves cancer cell lines, the cross design and its CSS scoring can be readily applied for ex-vivo drug screening, where the amount of patient-derived materials can be extremely limited and technically difficult to obtain due to culture constraints [23]. With the help of CSS and its visualization tools, drug combination discovery can be more quickly advance and may eventually lead to the validation of personalized drug combinations in clinical trials.

## Materials and Methods

### The cross drug combination design

We proposed a cross design to test the synergy and sensitivity of a drug pair by first introducing the concepts of background drug and foreground drug: background drug is the drug fixed at its IC_50_ concentration while foreground drug is added into the background drug with multiple concentrations. We allow that either drug in the pair to be the background drug, so that two vectors of dose mixtures will be produced and intersected at the IC_50_ concentrations (Fig 4). The dose-response curves for these two vectors will be determined using cell viability or toxicity assays, where the inhibition percentages can be calculated using negative and positive controls. Note that the cross design requires specifically the combinations at the IC_50_ concentrations, which need to be determined based on single drug sensitivity screening or prior knowledge.

**Fig 4.**
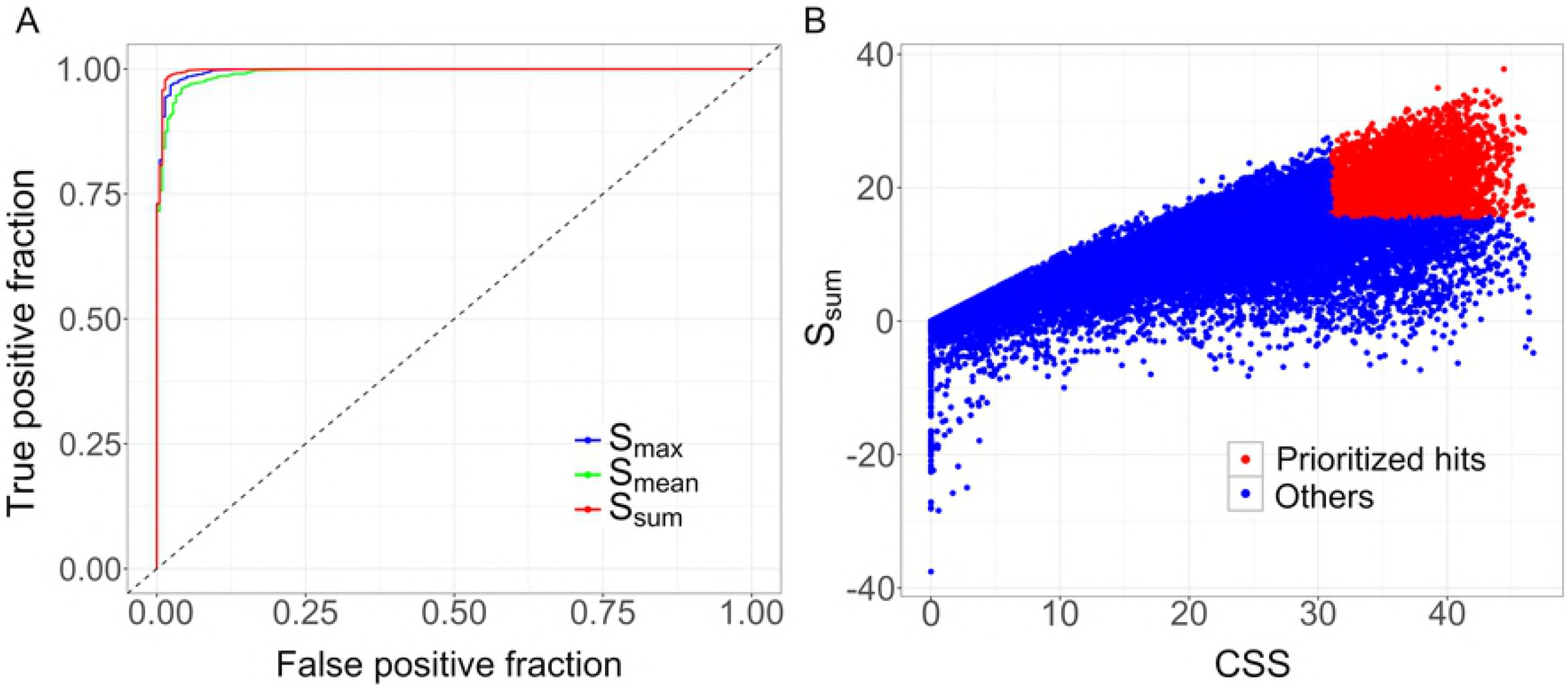
The cross design to determine the drug combination sensitivity score. Compared to the full-dose response matrix design (left panel), only the single row and single column that correspond to the IC_50_ concentrations of the two drugs were utilized for the calculation of CSS (middle panel). Either Drug1 or Drug2 can be considered as the background drug fixed at its IC_50_ concentration while the other is considered as the foreground drug with multiple doses being titrated. The resulting two dose-response curves will be summarized as the drug combination sensitivity score (CSS), from which a synergy score can also be calculated as the deviation from the expected value when there is no interaction.

### Determination of the CSS drug combination sensitivity scores

With the drug combination dose-response curves determined in the cross design, the CSS summarizes the area under the curve similar to the AUC and DSS (Drug Sensitivity Score) scoring approaches [24–25]. Namely, a four-parameter log-logistic function is used to fit the dose-response curve for a concentration *x* of the foreground drug according to:

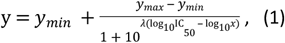

where *y_min_* and *y_max_* are the minimal and maximal inhibition (the bottom and top asymptotes of the curve,0 ≤ *y,y_min_,y_max_* ≤ 1; IC_50_ is the concentration of the foreground drug with which the drug combination reaches 50% inhibition of the cell growth; *λ* is the slope of the dose-response curve.

The dose-response curve (1) is transformed by substituting x with *x*’ = log_10_(*x*) as:

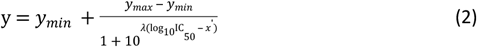

The area under the log_10_-scaled dose-response curve (AUC) is determined according to

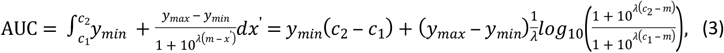

where [*c_1_ c_2_*] is the log_10_ concentration range for the foreground drug tested in the experiment, and m = log_10_(IC_50_).

The AUC is further normalized as the proportion of its theoretical upper bound according to:

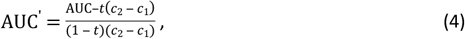

where *t* is the minimum inhibition level that is considered meaningful (by default it is fixed at 10%, assuming that the inhibition below 10% is experimental noise).

The CSS for the foreground drug is defined as a percentage and varies between 0 and 100:

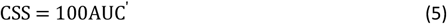

As there are two drug combination dose-response curves depending on which drug is fixed as the background drug, we refer to the results of Eq. (5) for either scenario as CSS_1_ and CSS_2_, and consider them as two samples that are generated from the same random variable. We take the average of CSS_1_ and CSS_2_ as the CSS for the drug pair, i.e.

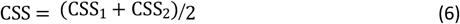

### The O’Neil drug combination data

Dose-response was measured as percentage of cell viability and retrieved from the supplementary material of [17], which includes 22,737 experiments for 583 drug pairs that involves 38 unique drugs in 39 cancer cell lines, representing 7 tissue types. At the first stage, single-drug screening was done using 8 concentrations to determine the IC_50_ concentration for each drug with six replicates. At the second stage, a 4 by 4 dose matrix was utilized to cover the span of IC_50_ concentrations for a drug pair with four replicates. To utilize the cross design, we picked up only the row and the column corresponding to the concentrations closest to the IC_50_ of the single drugs. These two vectors thus allowed the fitting of drug combination dose-response curves with which the CSS can be calculated. The cell viability percentage was first transformed to inhibition percentage according to:

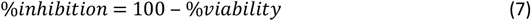

In our analysis, the average % inhibition of the four replicates was used to calculate the CSS. The robustness of the CSS values was assessed using the Pearson correlation across the four replicates.

### Predicting the CSS using machine learning approaches

With the CSS being determined for each drug combination, we sought to evaluate the prediction accuracy of multiple machine learning methods. We considered a drug combination as a combination of their targets and chemical fingerprints. We collected the known targets that have been experimentally validated for the 38 drugs from Drugbank [26] and ChEMBL [27]. Furthermore, we also utilized the Similarity Ensemble Approach (SEA) to predict additional secondary targets based on the chemical structures of the drugs [28]. The targets that were predicted with Z-score higher than 20, Tanimoto coefficient higher than 0.4 and P-value smaller than 0.01 were included. The MACCS fingerprints of the drugs were determined using the SMILES strings with the R package *rcdk* [29]. The resulting feature set for a drug combination altogether included 398 validated and predicted targets and 166 MACCS fingerprints (Table S1 and S2).

We compared three state-of-the-art machine learning methods for the CSS prediction: Elastic Net [30], Random Forests [31] and Support Vector Machines [32]. Elastic Net is a regularization and feature selection method that combines both ridge and lasso regression by including the L_1_ and L_2_ penalty terms. This method depends heavily on its penalty term that is regulated by hyper parameters α and *λ*. In our studies, α was selected from the interval [0.1, 1] and *λ* was chosen to minimize the difference between predicted and actual CSS scores. Random Forests is an ensemble learning method that constructs multiple decision trees. In our studies, we set the number of randomly selected predictors that is used at each split of the decision tree equal to the rounded down square root of the number of variables. For Support Vector Machines, the tuning parameters are the cost parameter *C* that sets the penalty for misclassification of a training point and a smoothing parameter *σ*, based on the accuracy of predictions in cross-validation.

We focused on the model performance for predicting new drug combinations within the same cell line, as the set of drug combinations in the training data did not overlap with that in the test data. For each cell line, we randomly sampled 70% of the drug combinations to train multiple machine learning models using 10-fold cross-validation. The optimized models were then used for predicting the CSS values for the remaining 30% of the novel drug combinations. The same processes were repeated 20 times by randomly splitting the training and test data. Four metrics including coefficient of determination (R2), root mean square error (RMSE), mean absolute error (MAE) and Pearson correlation (COR) were utilized for model comparison. To benchmark the performance of the machine learning methods, we utilized one randomly selected technical replicate as the prediction to obtain the upper limit of the performance. All the methods were implemented and evaluated using the R package *caret* [33].

### Determination of the CSS-based drug synergy scores

The advantage of CSS is that it allows a direct comparison of the sensitivity between a drug combination and its single drugs, and hence facilitates the quantification of drug synergy. The degree of synergy is often calculated as the deviation of the observed drug combination effect from the reference, which is defined as the expectation effect if the drugs are not interacting. We defined three variants of CSS-based synergy scores (termed as S scores) as:

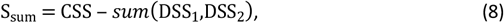

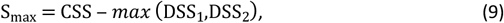

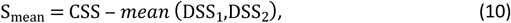

where the expectation was determined as a summary statistics of the normalized AUC based on the single drug dose-response curves (termed as DSS values as proposed in [25]). For a non-synergistic drug pair, the S score is expected to be zero. To evaluate the prediction accuracy of the CSS-based synergy scores, we defined a set of true synergistic and antagonistic drug combinations as the gold standard, which were determined using the full dose-response matrix data. We utilized the R package *synergyfinder* [34] to calculate multiple versions of synergy scores including the HSA (Highest single agency, [35]), the Bliss [36], the Loewe [37] and the ZIP synergy scores [38]. The principles of these four models were briefed as below:

Consider that drug 1 at concentration *x_1_* and drug 2 at concentration *x_2_* were combined to produce the inhibition effect of *y_c_*, while their respective single drug effects were *y*_1_(*x_1_*) and *y_2_*(*x_2_*).

The synergy score was calculated as the difference between *y_c_* and the expected effect *y_e_* if there is no synergy. Each synergy scoring took a different model for *y_e_*:

1. HSA: *y_e_* is the maximal single drug effect, defining

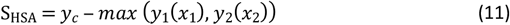
2. Bliss: *y_e_* is the expected effect of two drugs acting independently, defining

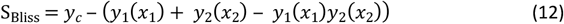
3. Loewe: *y_e_* is the expected effect of a drug combined with itself, defining

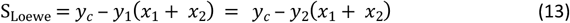
4. ZIP: *y_e_* is the expected effect of two drugs that do not potentiate each other, defining

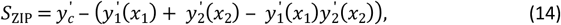

where 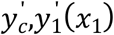 and 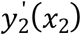 are the fitted values based on the full-dose response matrix for the combination and single drugs, respectively.

For each of the four models, the synergy scores were determined first for a given dose combination and then were averaged over the full dose-response matrix. With the four synergy scores determined for each drug combination, the true synergistic and antagonistic drug combinations are those with all the four synergy scores consistently higher than 5 and lower than 5, respectively. The aim was then to use the CSS-based synergy score which was determined by the cross design data to predict the ground truth determined by the full dose-response matrix design. The area under the ROC curve was used for evaluating how well the CSS-based synergy scores can predict the consensus drug combinations determined using the full dose-response matrix data.

## Acknowledgements

This work was supported by the European Research Council Starting Grant agreement No 716063 (DrugComb), Academy of Finland Grant agreement No. 317689 and Helsinki Institute of Life Sciences Research Fellow funding. A.M and W.W. are supported by the FIMM-EMBL PhD program scholarship. We thank the authors of the O’Neil study for making the drug combination data fully accessible.

## Supporting Information

Table S1. Drug target profiles including experimentally-validated primary and secondary targets, and SEA-predicted secondary targets for the 38 compounds [SupTable1.xlsx]

Table S2. The chemical information including MACCS fingerprint profiles for the 38 compounds [SupTable2.xlsx]

Table S3. The correlations of the CSS values obtained using the average of four viability replicates, with the CSS values obtained from the replicate separately [SupTable3.xlsx].

Table S4. CSS drug combination sensitivity scores and S synergy scores for each drug combination [SupTable4.xlsx]

Fig S1. The heatmap of the CSS1-CSS2 correlations sorted by drug combinations. Drug classes are shown in different colors at the edges of the heatmap.

Fig S2. Replicability of CSS values over the replicates. The line plot of the minimal and maximal values for the CSS replicates combined with CSS values over the standard deviation of the CSS replicates. Fig S3. The coefficient of variation (CV) for each drug in the single drug screens. The correlations between CSS1 and CSS2 for the drug combinations that involve a given drug were shown on top of each bar.

Fig S4. The correlation between the variable importance of TOP1MT and the average CSS for TOP1MT inhibitor for all the 39 cell lines. LNCAP is the only line which has a negative variable importance for TOP1MT.

